# A cell cycle kinase-phosphatase module restrains PI3K-Akt activity in an mTORC1-dependent manner

**DOI:** 10.1101/2020.11.26.399915

**Authors:** Belén Sanz-Castillo, Begoña Hurtado, Aicha El Bakkali, Dario Hermida, Beatriz Salvador-Barbero, Diego Martínez-Alonso, José González-Martínez, Clara Santiveri, Ramón Campos-Olivas, Pilar Ximénez, Javier Muñoz, Mónica Álvarez-Fernández, Marcos Malumbres

**Affiliations:** Cell Division and Cancer group, Spanish National Cancer Research Centre (CNIO), Madrid; Spectroscopy and Nuclear Magnetic Resonance Unit, Spanish National Cancer Research Centre (CNIO), Madrid; Proteomics Unit, Spanish National Cancer Research Centre (CNIO), Madrid

**Keywords:** AKT, ARPP19, cell cycle, ENSA, glucose homeostasis, MASTL, mTOR

## Abstract

The AKT-mTOR pathway is a central regulator of cell growth and metabolism. Upon sustained mTOR activity, AKT activity is attenuated by a feedback loop that restrains upstream signaling. However, how cells control the signals that limit AKT activity is not fully understood. Here we show that MASTL/Greatwall, a cell-cycle kinase that supports mitosis by phosphorylating the PP2A/B55 inhibitors ENSA/ARPP19, inhibits PI3K-AKT activity by sustaining mTORC1- and S6K1-dependent phosphorylation of IRS1 and GRB10. Genetic depletion of *MASTL* results in an inefficient feedback loop and AKT hyperactivity. These defects are rescued by expression of phospho-mimetic ENSA/ARPP19 or inhibition of PP2A/B55 phosphatases. MASTL is directly phosphorylated by mTORC1, thereby limiting the PP2A/B55-dependent dephosphorylation of IRS1 and GRB10 downstream of mTORC1. Downregulation of *MASTL* results in increased glucose uptake in vitro and increased glucose tolerance in adult mice, suggesting the relevance of the MASTL-PP2A/B55 kinase-phosphatase module in controlling AKT and maintaining metabolic homeostasis.

## Introduction

Cellular responses to growth factors such as insulin are transduced into the cell through multiple signaling cascades including the PI3K-AKT pathway ^1^. Upon stimulation, the PI3K-AKT pathway activates mechanistic target of rapamycin (mTOR) signaling through AKT-dependent phosphorylation of the tuberous sclerosis complex 2 (TSC2, tuberin) protein. Upon activation, the mTORC1 complex phosphorylates critical regulators of ribosomes and translation such as the ribosomal S6 kinases (S6K1 and S6K2) and the eukaryotic translation factor 4E (eIF4E)-binding protein 1 (4E-BP1), thus driving the synthesis of proteins required for cell growth and proliferation ^2-5^.

The mTORC1-S6K1 axis also mediates potent feedback loops that restrain upstream signaling through insulin/IGF receptor. S6K1 inhibits insulin receptor substrate-1 (IRS1) through a priming phosphorylation that sensitizes IRS1 to further inhibitory phosphorylation by other kinases ^6^. *TSC1/2*-deficient cells are characterized by insulin/IGF-1 resistance as a consequence of constitutive downregulation of IRS1 and IRS2 due to sustained mTOR activity ^6, 7^. Additional IRS1-independent feedbacks have been proposed, including the mTORC1-dependent phosphorylation of the growth factor receptor-bound protein-10 (GRB10), a negative regulator of insulin and IGF-1 signaling ^8, 9^. Pharmacological inhibition of mTOR relieves the feedback inhibition of AKT ^6, 7^, limiting the efficacy of therapeutic strategies aimed to prevent mTOR signaling ^10, 11^.

Although the involvement of phosphatases in this feedback has been far less studied, evidences suggest that the action of mTOR in the phosphorylation and degradation of IRS1 results from inhibition of PP2A ^12^. In fact, PP2A is able to dephosphorylate IRS1 ^13^ and inhibition of PP2A is sufficient to induce IRS1 degradation ^12^. The Serine/threonine kinase MASTL (also known as Greatwall) is a critical inhibitor of PP2A complexes during mitosis ^14-16^. MASTL is activated by cyclin-dependent kinase 1 (CDK1) complexes during mitotic entry leading to the phosphorylation of the endosulfines, ensosulfine α (ENSA) and the 19-kD cAMP-regulated phosphoprotein (ARPP19)^17, 18^. When phosphorylated, these small molecules function as specific inhibitors of phosphatase complexes in which PP2A activity is modulated by the B55 family of regulatory proteins (B55α, β, γ, δ). Loss of MASTL results in mitotic defects such as defective chromosome condensation and segregation errors in *Drosophila* and mammalian cells ^19-22^.

In this manuscript, we report a mitotic-independent function of the MASTL-ENSA/ARPP19-PP2A-B55 axis in modulating the response to glucose and the feedback loops induced by sustained mTOR-S6K1 activity. In conditions of nutrient excess and high mTOR signaling, MASTL inhibits PP2-B55 activity thereby preventing the dephosphorylation of the feedback targets IRS-1 and GRB10. These observations identify a new layer of control that interconnects a cell-cycle module with the negative feedbacks regulating the AKT-mTOR pathway, and suggest the possible use of MASTL inhibition to specifically limiting the effects of these mTOR-S6K1-mediated feedback loops.

## Results

### MASTL regulates AKT inhibition by the mTORC1-S6K1 axis

To study the effect of the MASTL-ENSA/ARPP19-PP2A-B55 pathway in nutrient signaling we first knocked down *MASTL* expression using specific short hairpin RNA (*shMASTL*) sequences in different human cell lines. These experiments were performed shortly after induction of protein knock down to avoid any interference with mitotic aberrations (Fig. S1a,b). Starvation of MDA-MB-231 cells for either growth factors or nutrients inhibited mTORC1 activity in control cells infected with a scrambled sequence, as shown by decreased phosphorylation of its substrates S6K1 and 4E-BP1 (Fig. 1a,b). However, *MASTL* knockdown resulted in increased phosphorylation of S6K1-T389, and increased phosphorylation of S6 and 4E-BP, especially in glucose-starved cells. These signals were accompanied by a significant increase in the inhibitory phosphorylation of TSC2 on T1462 (Fig. 1b), suggesting that *MASTL*-depleted cells are largely resistant to inhibit mTORC1 pathway in response to glucose deprivation.

**Figure 1.**
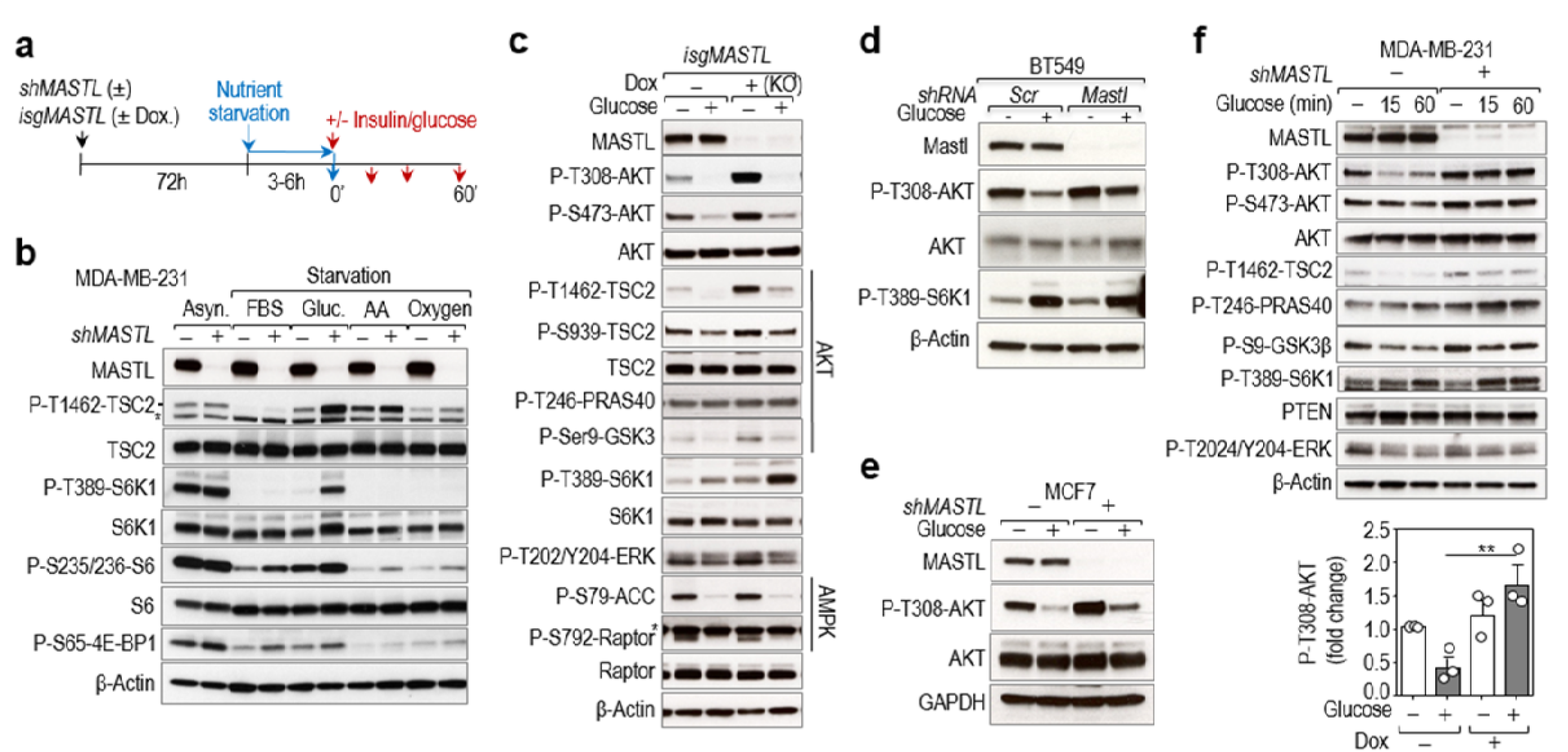
MASTL modulates the AKT-mTORC1 signaling pathway. **a)** Schematic representation of the protocol followed for depletion of MASTL in human cell lines and different nutrient starvations. *MASTL* was knocked down (RNA interference; *shMASTL*) or knocked out (inducible CRISPR/Cas9; *isgMASTL*) and cells were nutrient starved for 3-6 before analysis (time 0’). In case of re-stimulation with glucose or insulin, cells were analyzed 10-60 min after re-stimulation. **b)** Immunoblot analysis of asynchronous (Asyn.) MDA-MB-231 cultures or cells subjected to different starvations [elimination of fetal bovine serum (FBS), glucose (Gluc.), amino acids (AA) or oxygen] using the indicated antibodies in the absence (−) or presence (+) of shRNAs against *MASTL*. **c)** *MASTL* was ablated using the *isgMASTL* system. 72 h after Dox, cells were starved of glucose in the media in presence of 10% dFBS for 1 h and then re-stimulated with 25 mM of glucose for 10 min before recovery. Whole-cell lysates were blotted with the indicated antibodies. **d,e)** Immunoblot analysis with the indicated antibodies in BT549 and MCF7 cell lines, respectively. MASTL knockdown was performed by infection with shRNAs for MASTL (+) or Scramble (−) as a control. **f)** *MASTL* was depleted using shRNAs against *MASTL* (+) or scrambled shRNAs (−) as control. Cells were starved from glucose for 2 h and re-stimulated with 5 mM glucose for 15 min and 1 h. Total extracts were recovered and tested for the indicated antibodies. The graph shows the average of phospho-AKT T308 from three independent experiments (data are mean ± s.e.m; *p<0.01; 1-way ANOVA). In **b-f**, β-Actin or GAPDH were used as a loading control and * in panels b indicates unspecific bands.

We next switched to an inducible CRISPR system recently established (*isgMASTL*) in which Cas9 was transcriptionally induced by doxycycline (Dox) in the presence of specific small guide RNAs (sgRNAs) against human *MASTL* sequences (see STAR methods section for details). In the presence of doxycycline, *MASTL* was efficiently ablated resulting in increased S6K1 T389 and TSC2 T1462 phosphorylation upon glucose deprivation (Fig. 1c). In line with these observations, the TSC2-inhibitory kinase AKT was strongly activated in *MASTL*-null cells, as shown by increased phosphorylation on T308 and S473 residues. Other AKT substrates, like PRAS40, or GSK3, were also more phosphorylated in the absence of MASTL. No significant differences were observed in the phosphorylation of ERK1/2, another positive regulator of the mTORC1 pathway upstream of TSC2, or AMPK substrates such as ACC and Raptor, in these conditions. The effect of *MASTL* ablation on the AKT-TSC2-S6K1 axis was specific as the induction of Cas9 alone in control cells did not modify the phosphorylation status of the components of the AKT-S6K1 pathway (Fig. S1c). No major differences in AKT or S6K1 phosphorylation were observed in response to amino acids, indicating a specific role of MASTL in glucose signaling (Fig. S1d). The effect of *MASTL* knock down in the phosphorylation of AKT in response to glucose was confirmed in two additional independent cell lines, BT549 and MCF7 (Fig. 1d,e).

How glucose regulates AKT signaling is not well understood. However, it has been shown that chronic activation of mTORC1 by glucose results in decreased AKT phosphorylation ^23^. In a time course of glucose stimulation, control cells responded to this glucose-mediated feedback loop by inhibiting AKT activity. In contrast, *MASTL*-null cells failed to reduce AKT T308 phosphorylation, despite having enhanced mTORC1 activity as shown by phospho-S6K1 T389 (Fig. 1f).

We next analyzed the role of MASTL in the regulation of AKT activity following mTORC1/S6K1 activation by insulin in a time course experiment (Fig. 2a). PI3K/AKT mediated activation of mTORC1 is known to trigger a negative feedback loop ultimately leading to the attenuation of insulin signaling ^24-26^. Genetic ablation of *MASTL* did not modify the initial rise in AKT activity but prevented its downregulation after 30 min, despite having comparable levels of mTORC1 activity, resulting in sustained TSC2 and S6K1 phosphorylation (Fig. 2a). A similar effect was observed in non-transformed murine myoblasts (C2C12; Fig. 2b). Knockdown of *TSC2* in MDA-MB231 cells using specific shRNAs triggered AKT inhibition in control cells but not in MASTL-depleted cells, in which AKT activity was maintained (Fig. 2c). In line with these observations, acute mTORC1 inhibition by rapamycin treatment suppressed the effect of the negative feedback loop and boosted AKT phosphorylation (Fig. 2d). Lack of MASTL mimicked the effect of rapamycin on AKT phosphorylation (Fig. 2d), therefore suggesting a role for MASTL in the modulation of this feedback mechanism.

**Figure 2.**
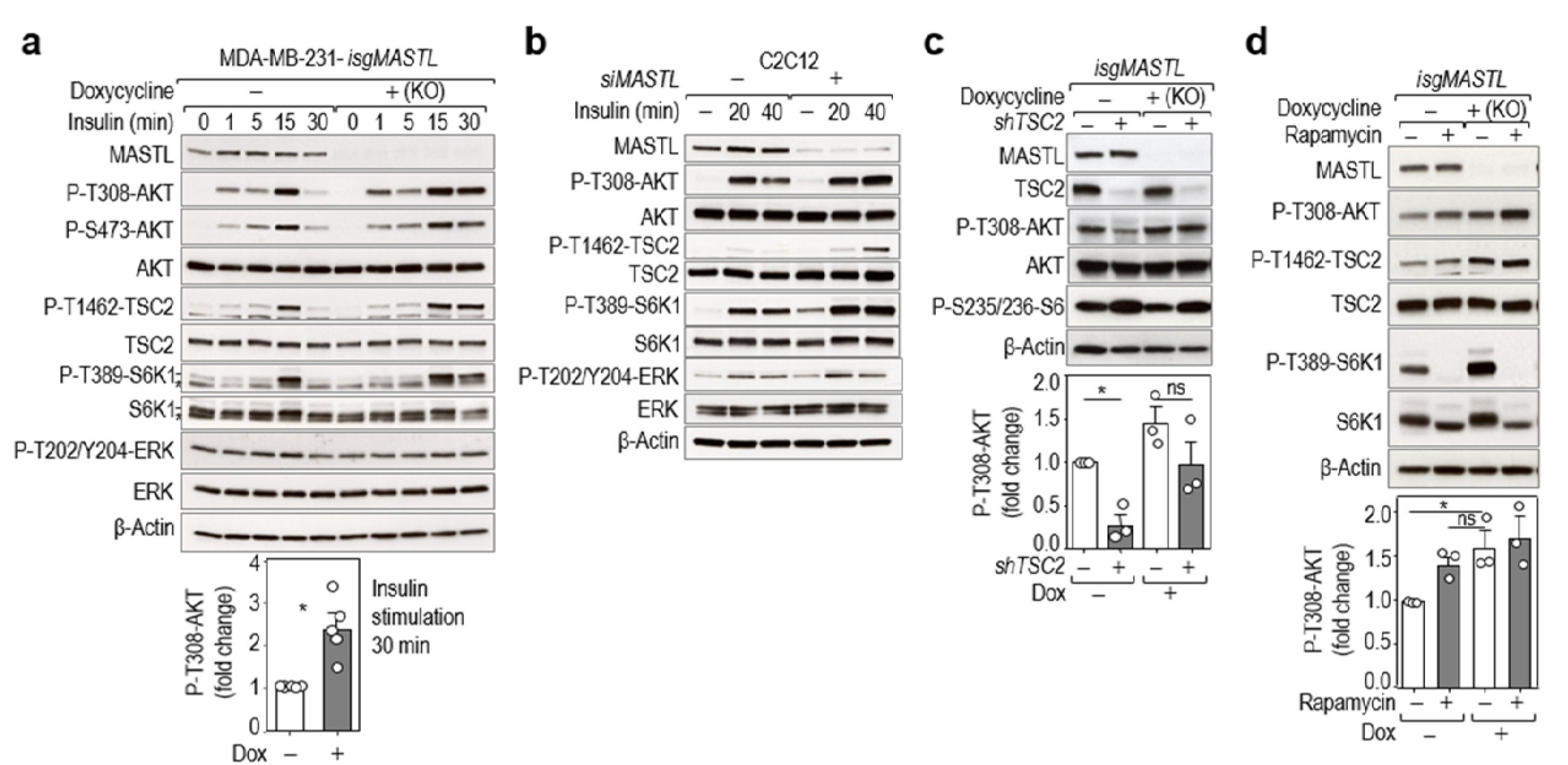
MASTL controls AKT phosphorylation in response to insulin signaling. **a)** Time course experiment at the indicated time points after 100 nM insulin stimulation in 6 h serum-starved control and *MASTL* knockout cells using the *isgMASTL* system. The histogram shows the quantification of phospho-AKT T308 after 30 min of insulin stimulation. n=5 independent experiments. * indicates unspecific bands. **b)** Mouse C2C12 myoblasts were depleted of Mastl using specific siRNAs. 48h after transfection, cells were placed in differentiation medium for 24h (2% horse serum), serum-starved for 5h and stimulated with insulin for the indicated periods of time. Whole-cell lysates were probed with the indicated antibodies. **c)** *TSC2* was knocked down in control or *MASTL*-null cells using specific (+) or Scrambled (−) shRNAs. Cells were starved and restimulated with insulin for 30 min. The histogram shows the quantification of phospho-AKT T308. n=3 independent experiments. **d)** Control (−) or *MASTL* null (+) cells were treated with 100 nM rapamycin or vehicle as control. Cells were starved for glucose for 1 h and re-stimulated with glucose for 15 min; rapamycin was added 15 min before glucose stimulation. The histogram shows the quantification of phospho-AKT T308. n=3 independent experiments. In **a-d**, β-Actin was used as a loading control. In panels **a,c,d,** data are mean ± s.e.m. ns, not significant. \**P*<0.05; two-sided Student’s *t*-test (**a**); 1-way ANOVA (**c,d**).

### MASTL depletion alters glucose metabolism in vitro and in vivo

The PI3K-AKT pathway is a central regulator of insulin-mediated glucose metabolism ^27^. Insulin stimulation following serum starvation increased GLUT4 translocation to the plasma membrane (PM), and *MASTL* knockdown further increased the percentage of cells positive for GLUT4 at the PM in this condition (Fig. 3a and Fig. S2a). Similar results were obtained upon glucose starvation and re-stimulation (Fig. S2b), further supporting a role for MASTL in the regulation of AKT-mediated GLUT4 translocation. In agreement with these data, depletion of MASTL led to a significant increase in glucose uptake compared with control cells (Fig. 3b). Analysis of the extracellular medium indicated that both glucose consumption and lactate production were increased upon *MASTL* silencing but to the same extent, so that the glycolytic index remained constant (Figure 3c,d). These results suggest that *MASTL* depletion does not induce a general metabolic switch, but rather a controlled and well-defined upregulation of glycolysis under conditions of feedback-mediated AKT regulation.

**Figure 3.**
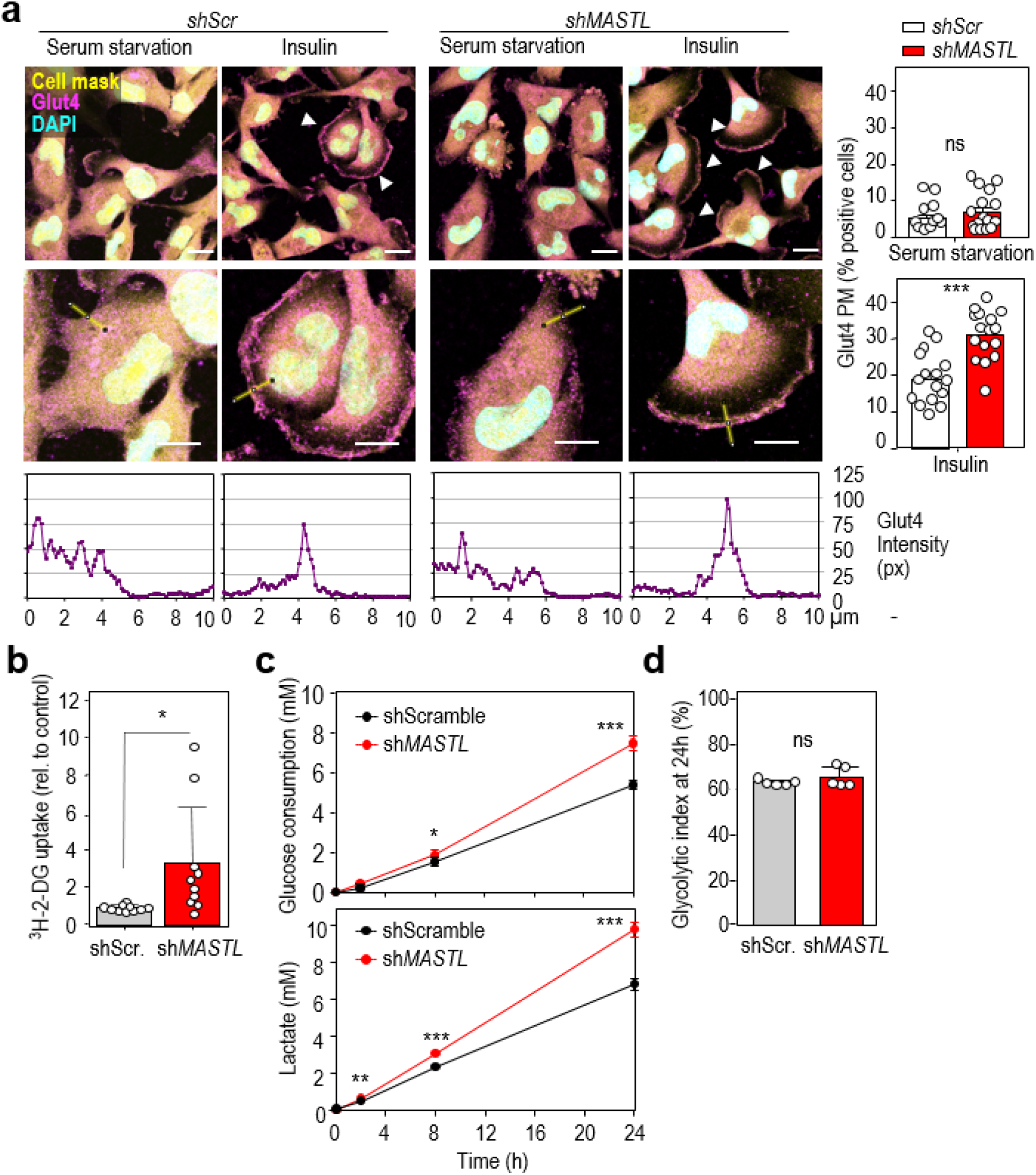
Metabolic alterations in MASTL-depleted cells. **a)** Immunofluorescence for GLUT4 (magenta) in MDA-MB-231 cells in conditions of serum starvation and insulin stimulation. DAPI (DNA) is in cyan and a cell mask is in yellow. Arrowheads indicate positive cells for GLUT4 enrichment at the plasma membrane (PM). Representative magnifications and line scan quantification of Glut4 intensity of the indicated representative sections are also shown. The histogram shows the quantification of the percentage of cells positive for GLUT4 translocation to the PM in 16 fields (800 cells) per condition. Scale bars, 10 μm. **b)** Cells were treated with specific shRNAs against *MASTL* (+) or scrambled sequences (−), and then serum-starved before incubation with 2-deoxy-D-[1-^3^H]-Glucose. Bars show mean + s.d. of 4 independent experiments with at least two replicates per experiment. \**P*<0.05, Student’s *t*-test. **c)** Control (shScr.) and *MASTL-depleted (shMASTL)* HepG2 depleted cells were incubated with 10 mM glucose and metabolites in the media were quantified by NMR at the indicated time points. Time course of glucose consumption and lactate production in cell media are indicated. The glycolytic index at 24 h (**d**) was calculated as −0.5*([Lac]/([Glc]0-[Glc]). Data in **c-d** are mean ± s.d. of 5 independent plates. ns, not significant; \**P*<0.05; \*\**P*<0.01, \*\*\**P*<0.001; two-sided unpaired Student’s *t*-test.

Mastl is expressed in pancreas, skeletal muscle, white adipose tissue (WAT) and liver mouse tissues at comparable levels to spleen, indicating that expression of this kinase is not restricted to proliferative tissues (Fig. 4a). Interestingly, Mastl expression was modulated in vivo in response to food intake in muscle, epididimal WAT and liver, in which *Mastl* mRNA levels increased in ad libitum fed mice compared to overnight fasted mice (Fig. 4b), further suggesting a non-mitotic role for Mastl in these tissues.

**Figure 4.**
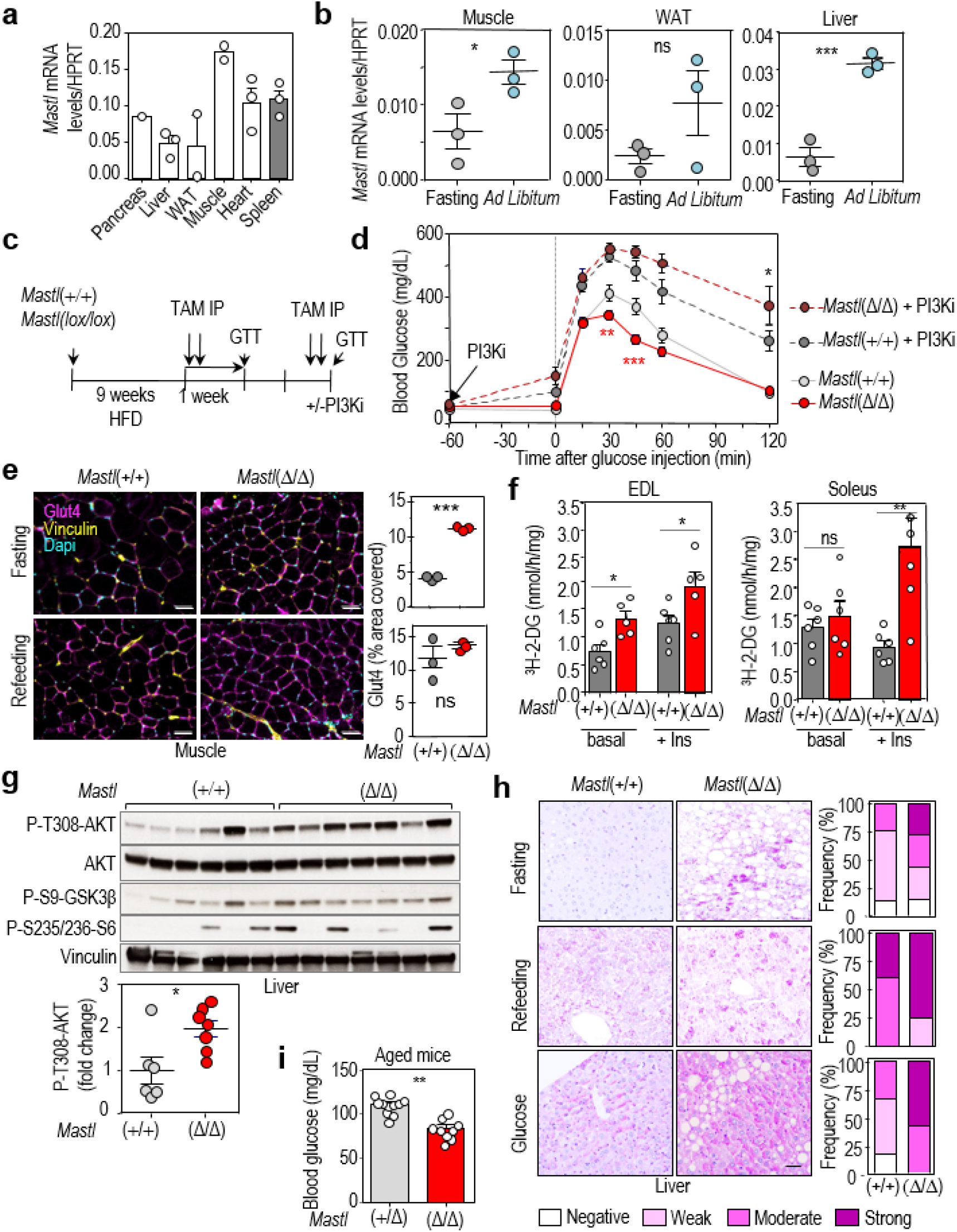
Mastl regulates glucose homeostasis in vivo. **a)** RT-qPCR analysis of *Mastl* transcripts in different tissues from *ad libitum* mice. The graph represents the average of normalized Mastl level from 3 mice (except in pancreas n=1). **b)** Relative levels of *Mastl* transcripts (normalized to *Hprt* mRNA levels) in muscle, epididymal white adipose tissue (WAT) and liver from wild-type mice after fasting for 16 h or fed *ad libitum*. **c)** Schematic representation of the protocol followed for the glucose tolerance test (GTT). TAM IP, tamoxifen intraperitoneal injection. **d)** Glucose tolerance test in *Mastl*(+/+) (n=6) and *Mastl*(Δ/Δ) (n=11) mice. 8-12-week-old male mice were fed with high fat diet (HFD; 60% fat) for 9 weeks before treated with tamoxifen to induce Mastl deletion. Treatment of *Mastl*(+/+) (n=5) and *Mastl*(Δ/Δ) (n=10) mice with the PI3K inhibitor ETP46992 (PI3Ki) was performed 1 h before glucose injection. **e)** Glut4 staining (magenta) in skeletal muscle sections. Vinculin (membrane) and DAPI (DNA) staining is shown in yellow and cyan, respectively. Mice were fasted overnight for 16 h, and re-fed for 2 h before sample collection. Quantification of the area covered by Glu4 staining [n=3 *Mastl*(+/+) and 3 *Mastl*(Δ/Δ)] is shown. Scale bar, 50 μm. **f)** Ex-vivo glucose uptake in EDL and soleus muscles from *Mastl*(+/+) (n=6) and *Mastl*(Δ/Δ) (n=5) mice, in basal conditions and upon insulin stimulation (100 nM, 20 min). Mice were fasted overnight before muscle isolation, and data were normalized to muscle weight. **g)** Immunoblot with the indicated antibodies in liver tissues from *Mastl*(+/+) (n=6) and *Mastl*(Δ/Δ) (n=7) mice. Mice were fasted overnight for 16 h, injected intraperitoneally with glucose (2 g/kg body), and sacrificed 30 min later for sample collection. Quantification of the relative fold change signal of phospho-AKT T308 (normalized to total AKT level). n=6 *Mastl*(+/+) and 7 *Mastl*(Δ/Δ) mice. **h)** Representative images of Periodic acid–Schiff (PAS) staining in liver sections from *Mastl*(+/+) and *Mastl*(Δ/Δ) mice. Mice were fasted overnight for 16 h, injected intraperitoneally with glucose (2 g/kg body) or re-fed for 2h and sacrificed 30 min later for sample collection. Bars show the relative frequency of PAS staining intensity according to the PAS score shown in Supplementary Fig. 4 (n=3 mice/ genotype). Chi-square test (\*\*\**P*<0.001). **i)** Blood glucose levels in *ad libitum Mastl*(Δ/Δ) (n=12) and *Mastl*(+/Δ) (n=11) mice. Mice were treated with tamoxifen diet to induce Mastl depletion and blood glucose concentration was determined 1-month later in the morning of ad libitum fed mice. In **a,b,d-g,i** data are mean ± s.e.m. ns, not significant; \**P*<0.05; \*\**P*<0.01; \*\*\**P*<0.001; 2-way ANOVA (**d**) or two-sided Student’s *t*-test (**b,e-g, i**).

We directly tested the relevance of Mastl in glucose metabolism in vivo by using a Mastl conditional knockout model in which the murine *Mastl* gene can be excised by a tamoxifen-induced Cre recombinase ^21^. Young (8-12-week-old) control and *Mastl*(lox/lox) knockout mice were fed with high fat diet (HFD, 60% of calories derived from fat, ~4 kcal/kg) for 9 weeks before the induction of Mastl deletion by tamoxifen injection (Fig. 4c). To avoid any interference with the potential proliferative defects induced by Mastl deletion, acute deletion was achieved in adult mice by intraperitoneal injection just before the metabolic assays. In these conditions, tamoxifen treatment and concomitant loss of Mastl did not affect the body weight of *Mastl*(Δ/Δ) compared to control *Mastl*(+/+) mice (Fig. S3a), and no significant defects were observed in highly proliferative tissues such as intestine or in weight gain (Fig. S3a,b). Although no statistical differences were observed in an insulin tolerance test (Fig. S3c), *Mastl*(Δ/Δ) mice displayed better glucose clearance (Fig. 4d), Importantly, acute treatment with an orally bioavailable specific PI3K inhibitor known to inhibit AKT activity ^28^ completely rescued the observed differences in glucose clearance (Fig. 4d). Of note, the effect of the inhibitor was even more pronounced in *Mastl*(Δ/Δ) mice, who reached the highest and most sustained levels of hyperglycemia, compared to control mice (Fig. 4d). These data also suggest that the enhanced glucose tolerance upon *Mastl* ablation might be a consequence of increased PI3K/AKT activity in *Mastl*(Δ/Δ) tissues responsible of glucose uptake, such as muscle, WAT or liver.

In agreement with these observations, expression of Glut4 at the plasma membrane (PM) increased in *Mastl*(Δ/Δ) muscles, especially in fasting conditions where mTORC1 pathway is usually downregulated in control tissues, suggesting mTORC1 and Akt hyperactivity in the absence of Mastl (Fig. 4e). Since most insulin-mediated glucose uptake occurs in skeletal muscle, isolated muscles derived from Mastl-deficient mice were assessed for their capacity to uptake ^3^H-2-deoxy-D-glucose. As shown in Fig. 4f, the rate of glucose uptake was significantly increased in *Mastl*(Δ/Δ) extensor digitorum longus (EDL) muscles in basal conditions, and even further upon insulin stimulation in both EDL and soleus muscles, representing fast- and slow-twitch muscles, respectively (Fig. 4f).

Phosphorylation of AKT was increased in response to glucose stimulation in *Mastl*(Δ/Δ) livers (Fig. 4g). AKT signaling in the liver promotes the conversion of glucose-6-phosphate into glycogen. Accordingly, liver sections from *Mastl*(Δ/Δ) mice showed a significant increase in periodic acid shiff (PAS) staining, which detects glycogen and other polysaccharides, during fasting and also upon refeeding or glucose stimulation (Fig. 4h). These data are consistent with alleviation of mTORC1-dependent negative feedback mechanisms affecting upstream insulin signaling in these muscles.

Finally, 1-year-old *Mastl*(Δ/Δ) mice fed with chow diet ad libitum had a significantly reduced glycaemia compared to control *Mastl*(+/Δ) littermates (Fig. 4i), suggesting better glucose clearance in aged Mastl-deficient mice. All together, these findings suggest that Mastl ablation improves glucose tolerance upon a stressful condition such as HFD, as well as basal glycaemia in old mice.

### MASTL controls AKT activity in an ENSA/ARPP19-PP2A-B55 dependent manner

We next tested whether MASTL could regulate AKT-mTORC1 signaling through the ENSA/ARPP19-PP2A pathway. As shown in Fig. 5a, knockdown of *ENSA* and *ARPP19* mimicked the effect of *MASTL* ablation, and resulted in increased AKT phosphorylation thereby preventing the effect of glucose in the inactivation of AKT by the feedback loop. In addition, expression of *ENSA* and *ARPP19* cDNAs with mutations that mimic MASTL-dependent phosphorylation (ENSA-S67D/ARPP19-S62D) restored the downregulation of phospho-AKT after MASTL ablation in conditions of glucose readdition (Fig. 5b). The involvement of PP2A was then tested in similar assays using PP2A phosphatase inhibitors which we would expect to rescue the effect of MASTL inhibition. Although these inhibitors are not specific for PP2A/B55 complexes, treatment of cells with the general PP2A inhibitors fostriecin or okadaic acid led to decreased levels of AKT phosphorylation on T308, partially restoring AKT inhibition in *MASTL-null* cells (Fig. 5c). Altogether these data indicate that the effect of MASTL on the regulation of AKT occurs through the ENSA/ARPP19-mediated inhibition of PP2A.

**Figure 5.**
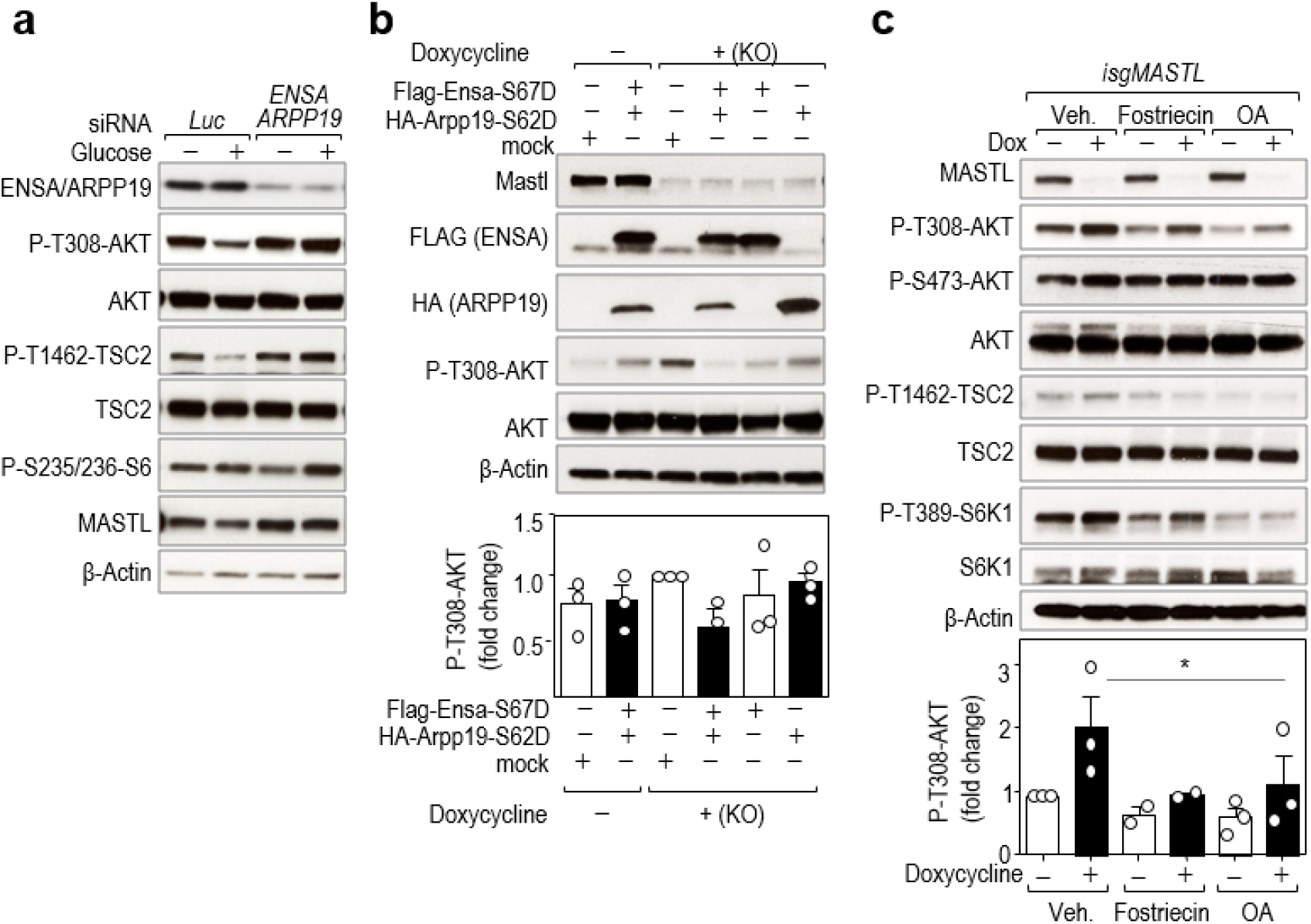
MASTL mediates mTORC1/S6K1-mediated AKT inhibition through the ENSA/ARPP19-PP2A/B55 pathway. **a)** Co-depletion of *ENSA* and *ARPP19* proteins in the MDA-MB-231 cell line using specific siRNAs or control siRNAs against luciferase (*Luc*). 72 h after transfection cells were starved for glucose for 1 h and re-stimulated with glucose for 15 min. Total cell extracts were collected and analyzed for the indicated antibodies. **b)** *isgMASTL* cells were previously infected with lentiviral supernatants expressing FLAG- and HA-tagged versions of ENSA and ARPP19 phospho-mimetic mutants (S67D and S62D) or GFP as a control. Cells were treated with Dox to induce *MASTL* CRISPR/Cas9-dependent knock-out, and 72 h later cells were treated as specified in **a)**. **c)** Control (Dox −) and *MASTL* knockout (Dox +) cells were treated with 5 μM fostriecin, 50 nM okadaic acid (OA) or vehicle (Veh) 10 min before insulin stimulation for 30 min. Total lysates were analyzed for the indicated antibodies. Histograms in **b, c,** represents the quantification of phospho-AKT T308 from 3 independent experiments. Data are mean ± s.e.m. \**P*<0.05; paired Student’s *t*-test.

### The MASTL-PP2A/B55 axis modulates phosphorylation of the feedback targets IRS1 and GRB10

AKT signaling has been shown to be modulated by PP2A complexes ^29^. However, the fact that MASTL inhibition and, as a consequence, PP2A/B55 activation leads to increased phosphorylation of AKT indicates that the effect we observe on the feedback loop is not mediated by direct dephosphorylation of AKT by PP2A/B55. Feedback inhibition of the PI3K/AKT pathway by mTORC1 involves mTORC1/S6K1-dependent phosphorylation of the insulin receptor substrates, IRS1 and IRS2, as well as the adaptor protein GRB10. Whereas phosphorylation of IRS1 results in reduced protein levels as a consequence of ubiquitin-dependent degradation ^30, 31^, GRB10 phosphorylation prevents proteosomal-mediated degradation thereby correlating with increased protein levels ^8, 32^. Interestingly, IRS1 stability is also regulated by PP2A complexes ^12^, although the precise mechanism remains unknown. As shown in Fig. 6a, knockdown of B55α and B55δ, the two ubiquitous B55 isoforms, led to increased phosphorylation of IRS1, mostly on S616, a phosphoresidue that negatively regulates IRS1 function. Whether phosphorylation of S312, another residue involved in the mTOR-mediated feedback inhibition of AKT, is also regulated by PP2A/B55 is not clear, given the very low signal obtained with this antibody. Phosphorylation of GRB10 on S476 was also increased upon B55α,δ depletion, and, as expected, these phosphorylations correlated with decreased AKT activation in B55α,δ-depleted cells (Fig. 6a). On the contrary, ablation of *MASTL* prevented the phosphorylation of both IRS1 S616 and GRB10 S476, suggesting defective feedback signaling, and resulted in increased IRS1 and reduced GRB10 total protein levels (Fig. 6b). Depletion of B55 subunits in *MASTL*-null cells partially rescued the phosphorylation status of IRS1 and GRB10, and the levels of AKT activity, although these assays were limited by technical difficulties in the efficient elimination of these B55 proteins (Fig. 6c).

**Figure 6.**
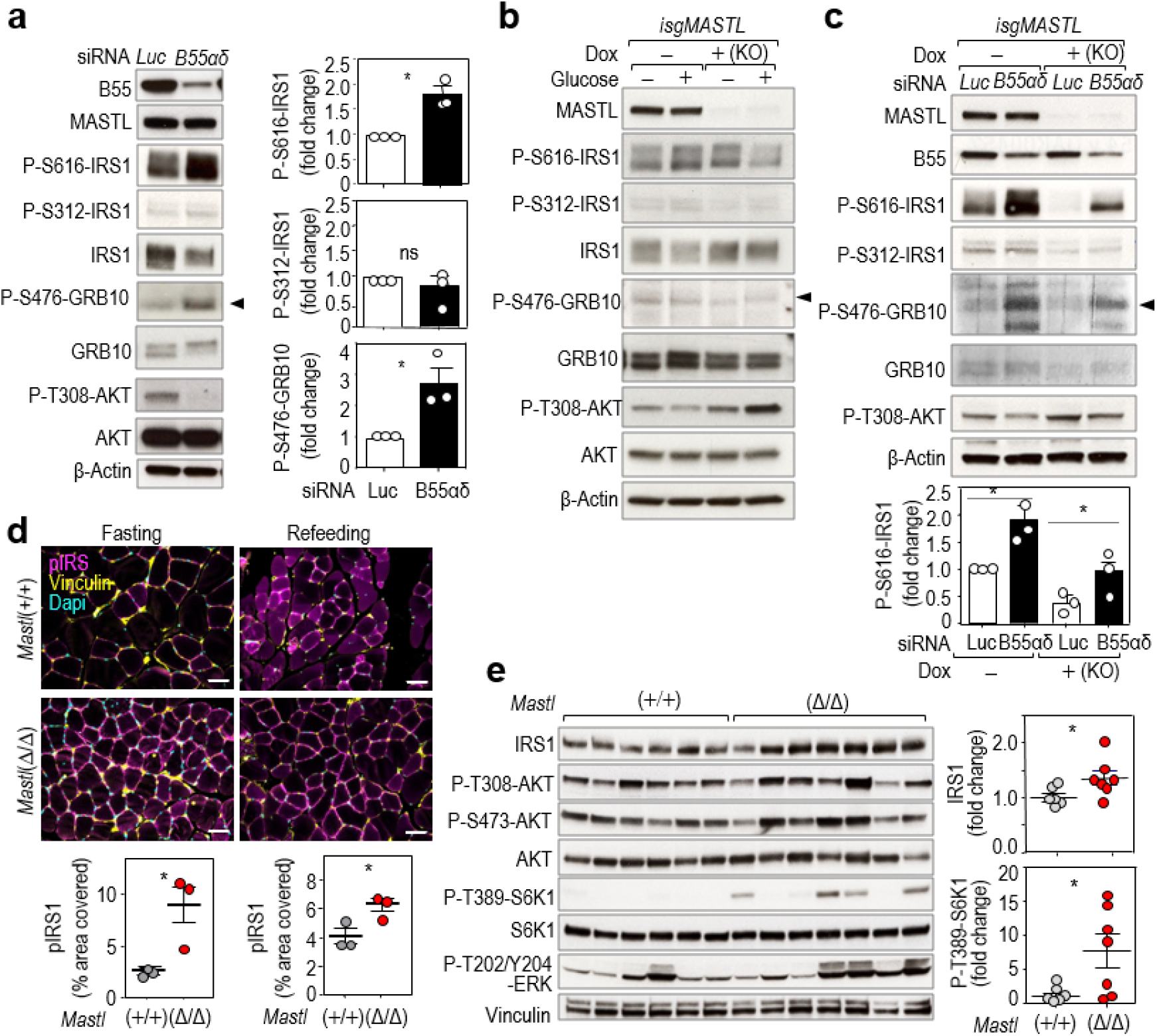
The MASTL-PP2A/B55 pathway controls the feedback loop at the level of adaptor proteins. **a)** MDA-MB-231 cells were co-transduced with specific siRNAs against the *PPP2R2A* (B55α) and *PPP2R2D* (B55δ) transcripts (B55αδ) or against luciferase (*Luc*) as a control. Asynchronous cells were collected and blotted for the indicated antibodies. The histograms represent the quantifications of the specified phospho-residues. n=3 independent experiments. **b)** Immunodetection of the indicated antigens in control (−) and *MASTL* knockout (+) cells starved from glucose and restimulated with glucose for 15 min. **c)** siRNA-mediated genetic depletion of B55alpha and delta subunits (*B55αδ*) in control (Dox −) and *MASTL-null* (Dox +) cells. The phosphorylation status of the different proteins was scored in conditions of glucose readdition. n=3 independent experiments. In **a,b,c** the phospho-S476 GRB10 signal is indicated by arrowheads. **d)** Phosphorylated IRS1 (S616) staining (magenta) in skeletal muscle sections. Vinculin (membrane) and DAPI (DNA) staining is shown in yellow and cyan, respectively. Mice were fasted overnight for 16 h, and re-fed for 2h before sample collection. Histograms show the quantification of the phospho-IRS1 mean intensity [n=3 *Mastl*(+/+) and 3 *Mastl*(Δ/Δ)]. Scale bar, 50 μm. **e)** Immunoblot with the indicated antibodies in skeletal muscle (gastrocnemius) extracts. Mice were fasted overnight for 16 h, injected intraperitoneally with glucose (2 g/kg body), and sacrificed 30 min later for sample collection. Quantification of the relative fold change signal of total IRS1, and phospho-S6K1 T389 (normalized to total S6K1 level; n=6 *Mastl*(+/+) and 7 *Mastl*(Δ/Δ)). β-Actin or vinculin were used as a loading control. In **a,c-e,** data are mean ± s.e.m. ns, not-significant; \**P*<0.05; one-way ANOVA (**c**) or two-sided Student’s *t*-test (**a,d,e**).

To further confirm these results in vivo, we analyzed the status of IRS in Mastl-deficient muscle samples. Muscle sections from *Mastl*(Δ/Δ) mice showed a significant increase in IRS both in fasting and refeeding conditions (Fig. 6d), and total IRS1 levels were higher in Mastl-*null* tissues compared to control ones, in agreement with a deficient feedback loop due to Mastl deletion (Fig. 6d).

### Control of MASTL activity by the mTORC1-S6K1 axis

As discussed above, glucose re-addition following glucose starvation triggers mTORC1-dependent AKT inhibition. In these conditions, phosphorylation of MASTL in one of the potential T-loop activation sites (T194) was increased (Fig. 7a). In agreement with these data, phosphorylation of MASTL on T194 was also increased in *TSC2*-null cells, and was prevented by the mTOR inhibitors rapamycin and torin (Fig. 7b). *In vitro* kinase assays using GFP-MASTL immunocomplexes showed that MASTL activity over recombinant ARPP19 increased upon *TSC2* knockout, and was partially prevented upon rapamycin and torin treatment (Fig. S4a). These data suggested that MASTL is under the control of the mTORC1-S6K1 axis in a mitosis independent manner. In fact, both kinases were able to phosphorylate MASTL in assays with recombinant proteins (Fig. 7c). Mass-spectrometry analysis identified T710, S716/T718, T722 and S878 as possible sites phosphorylated by mTORC1. Residues T710, T716/T718, T722 were also phosphorylated by S6K1 although to a lesser extent. Among the four mTORC1/S6K potential sites, S878 contains a hydrophobic residue in position +1, fitting with the consensus motif for mTORC1 substrates ^33, 34^. Interestingly, S878 was exclusively phosphorylated by mTORC1 and to a much higher extent compared to the other identified sites (150 fold versus less than 10 fold for T710, T716/718 or T722; Fig. 7d). These four sites are located in the C-terminal part of MASTL and presented high conservation among species (Fig. 7e).

**Figure 7.**
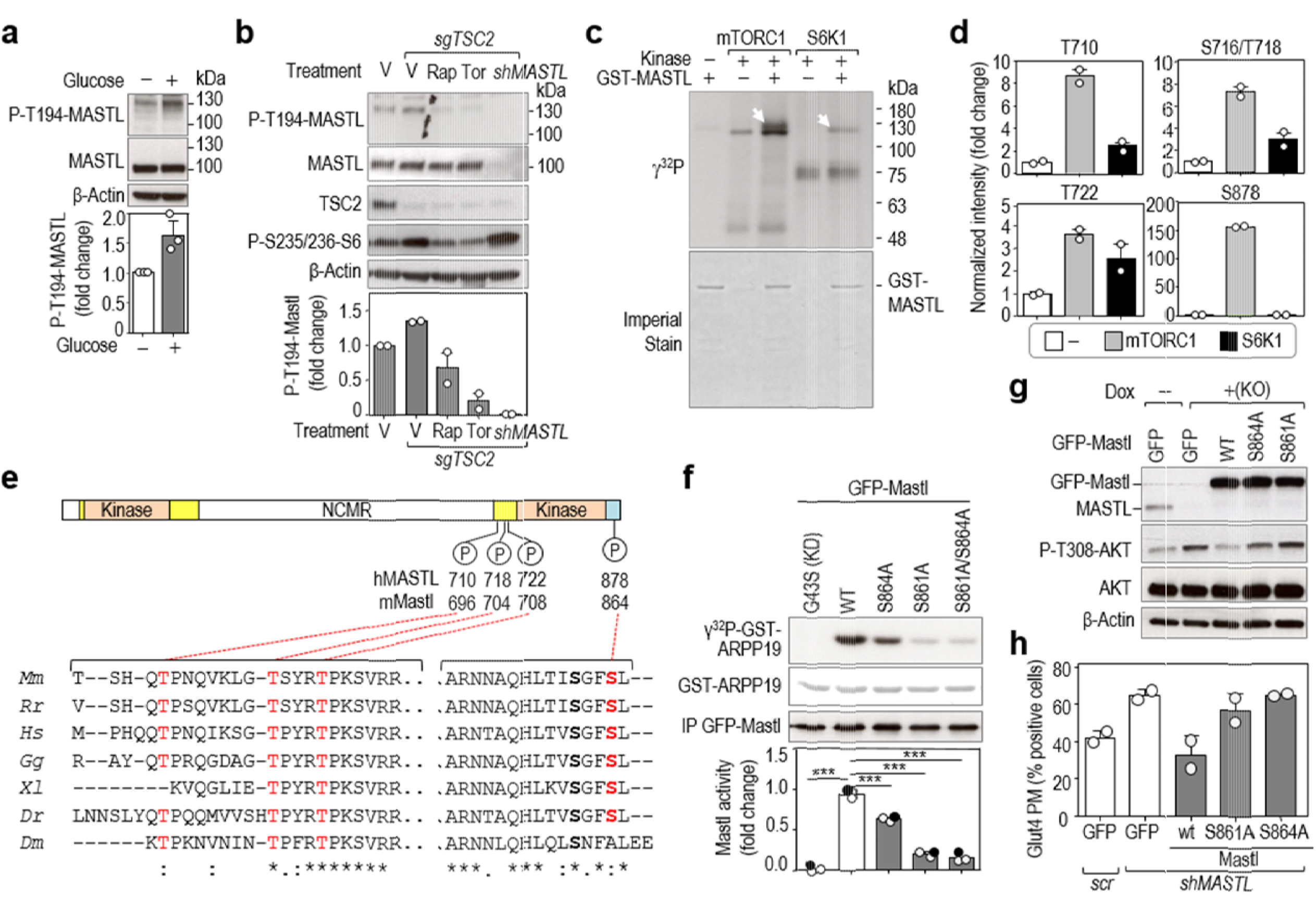
The mTORC1/S6K1 pathway regulates MASTL kinase activity. **a)** MDA-MB-231 cells were starved for 1 h and re-stimulated with glucose for 15 min and T194 phosphorylation was tested with a specific antibody. The histogram shows the quantification of phospho-MASTL T194 (n=3 independent experiments). **b)** Cas9-expressing control (−) cells or cells with CRIPSR/Cas9-mediated *TSC2* knockout (*sgTSC2*) were treated with rapamycin (Rap), Torin1 (Tor), or vehicle as control (V). *MASTL*-knockdown cells (*shMASTL*) were used as an additional control of the specificity of the antibody. Inhibitors were added 15 min before glucose re-stimulation for additional 15 min and total cell lysates were harvested for immunoblot. The histogram shows the quantification of phospho-MASTL T194 (n=2 independent experiments). **c)** In vitro kinase assay using purified GST-human MASTL as a substrate, and recombinant mTORC1 and S6K1 kinases. Arrowheads show the phosphorylation of MASTL in the presence of mTORC1 or S6K1. Note the appearance of other phosphorylation signals in mTORC1 and S6K1 reactions, independently of the presence of MASTL, which correspond to autophosphorylation of these kinases. Imperial staining shows the amount of GST-MASTL protein used as a substrate in each reaction. **d)** Mass spectrometry analysis of samples treated as in panel **c)**. The histograms represent the intensity of mTORC1-S6K1-regulated phospho-sites on human MASTL. Data (mean ± s.d.; n=2 technical replicates) were normalized to the total levels of MASTL and plotted as relative values to the condition of MASTL alone. **e)** Schematic representation of MASTL protein structure showing the phosphorylation sites identified in the in vitro kinase assay shown in c). Orange boxes indicate the conserved kinase domains. Yellow boxes indicate additional sequences conserved in Mastl in different species. The C-tail domain is shown in blue. The alignment of the amino acid sequence regions surrounding those phosphosites across different species is shown below. Phosphoresidues are highlighted in red colour. *Mm, Mus musculus; Rr, Rattus rattus; Hs, Homo sapiens; Gg, Gallus gallus; Xl, Xenopus laevis; Dr, Danio rerio; Dm, Drosophila melanogaster*. **f)** In vitro kinase activity of Mastl mutant isoforms. HEK-293T cells were transiently transfected with the indicated GFP fusions of mouse Mastl mutants. Anti-GFP immunocomplexes were subjected to a kinase reaction using recombinant ARPP19 (white circles) or ENSA (filled circles) as a substrate. The graph shows the relative catalytic activity of the indicated mutants. Kinase activity of GFP-mMastl wild-type was set as 1 (n=3 independent experiments). **g)** *isgMASTL* MDA-MB-231 cells were transduced with the indicated Mastl mutants or GFP alone as a control, and treated with doxycycline (Dox) to induce *MASTL* deletion. Cells were collected after glucose stimulation, and total lysates blotted for the indicated antibodies. **h)** MDA-MB-231 cells were transduced with the indicated Mastl mutants or GFP alone as a control, and infected with scramble or sh*MASTL* to deplete endogenous *MASTL*. Cells were fixed upon insulin stimulation for Glut4 staining. Quantification of the percentage of cells positive for GLUT4 at the plasma membrane in each condition is shown. (n=2 independent experiments) Data are mean + s.d. (**d**) or ± s.e.m (**a**, **b**, **f and h**). \**P*<0.05; \*\**P*<0.01; \*\*\**P* <0.001; two-sided Student’s *t*-test (**a**) or one-way ANOVA (**b, f**).

MASTL belongs to the AGC kinase family but it differs from other family members in a 500 aa-long insertion of unknown role in the activation loop that divides the kinase domain into two separate N-terminal and C-terminal fragments (Fig. 7e). The C-terminal tail (known as the turn motif in PKA or the tail/Z site in growth factor-stimulated AGC kinases) is a critical region whose phosphorylation is required for conformational changes and kinase activity. In fact, autophosphorylation at S875 is considered a major requirement for MASTL activation, which requires some priming phosphorylations ^35, 36^ and S878, the main mTOR phosphorylation site, locates next to it in the C-terminal tail (Fig. 7d and Fig. S4c). Independent proteomic screens indicate that both S875 and S878 are phosphorylated in mammalian and *Xenopus* mitotic extracts (www.phosphosite.org). Phosphorylation of S878 has also been found in interphase with high occupancy, and proposed as one of the potential priming phosphosites for MASTL activation ^36^. We actually detected phosphorylation of MASTL S878 in non-mitotic (G1) cells in response to growth factors (Fig. S4b). Mutation of the homologous residue in *Xenopus* (S886) results in a partial inactivation of the protein in mitotic extracts [about 60% kinase activity on its substrates; ^35^]. We detected a similar reduction in phosphorylation of recombinant ARPP19 or ENSA (about 60% compared to wild-type Mastl) when the homologue mouse residue (S864) was mutated to Alanine and immunopurified from interphasic cells (Fig. 7f). Mutation of the autophosphorylation site (mouse S861A equivalent to human S875) almost completely abolished Mastl kinase activity, and we did not detect any further reduction in catalytic activity when both mutants were combined, suggesting that phosphorylation of S864 might work as a priming phosphorylation site for mTORC1-dependent MASTL activation (Fig. 7f).

To evaluate the relevance of the phosphorylation of MASTL by mTOR in the feedback in response to glucose stimulation, we performed reconstitution assays using *MASTL* knock-out cells expressing GFP-fusions with the mouse Mastl mutants resistant to the sgRNAs against human *MASTL*. Whereas the wild-type version of Mastl was able to restore the phosphorylation of AKT in T308, mutation of Mastl in the mTOR site (S864A) was not able to reduce AKT activity to the level of control cells (Fig. 7g). Similarly, this Mastl phospho-mutant also failed to rescue the increment in Glut4 at PM observed in Mastl-deficient cells (Fig. 7h). Collectively, these data suggest that mTORC1 might modulate the kinase activity of MASTL, through the phosphorylation of its C-terminal region, thereby inhibiting PP2A-B55 in order to restrain AKT activity in response to nutrients.

## Discussion

The presence of negative feedback loops (Fig. S4d) that attenuate AKT activity upon sustained mTORC1-S6K signaling reflects the need for fine-tuning regulation of the AKT-mTOR axis at different levels. Mechanistically, suppression of AKT depends on the mTORC1- and S6K1-dependent phosphorylation of IRS1, IRS2, and GRB10 ^6-9^, as well as non-cell autonomous effects such as the mTORC1-dependent secretion of IGF binding protein 5 (IGFBP5) which blocks IGFR-I activation in the extracellular space ^37^.

The participation of cellular phosphatases in these feedback loops has been far less studied. PP2A inhibition has been previously shown to increase IRS1 phosphorylation inducing its degradation ^12, 13^. Because of the multiplicity of regulatory subunits, the involvement of specific PP2A complexes in different molecular pathways is not well established. Our data indicate that PP2A complexes containing the B55 family of regulatory subunits modulate the phosphorylation status of IRS1 and GRB10, and are inhibited by MASTL-ENSA/ARPP19 during feedback signaling.

A few studies have previously linked MASTL with AKT signaling. MASTL overexpression in MCF10-A cells is accompanied by increased phosphorylation of GRB10 on S104, and of IRS1 on S312 (two residues also involved in the AKT-mTORC1 negative feedback loop), and decreased phosphorylation of TSC2 on S939, an AKT site involved in the regulation of mTORC1 activity ^38^. In contrast, exogenous expression of MASTL may also induce AKT hyperactivation in tumor cells ^39^. Intriguingly, this effect is ENSA-PP2A/B55 independent and affects phosphorylation of AKT on S473, but not T308, perhaps suggesting that the regulation of mTORC2 and/or PHLPP1, the main kinase/phosphatase responsible for AKT S473 phosphorylation may be PP2A independent. It is therefore possible that MASTL may influence AKT signaling differentially in cycling ^39^ or starved/G0 cells as suggested by our data. Interestingly, AKT may lead to MASTL phosphorylation and activation specifically during mitosis ^40^, although whether this is mediated by direct phosphorylation and whether these data have any relevance in vivo in a physiological context has not been addressed so far.

The MASTL-PP2A kinase-phosphatase module is highly conserved across organisms although its physiological functions have varied through evolution. In the budding yeast, the MASTL-Endosulfine (RIM15-IGO1/2) axis is required for G1 arrest and entry into quiescence in response to nutritional deprivation by inducing the expression of a particular transcriptional response ^41-44^. In fission yeast, TORC1 activity leads to inhibition of the MASTL orthologue Ppk18 in the presence of nutrients. In that situation, cyclin B-Cdk1 complexes are inhibited in a PP2A-depednent manner delaying mitotic entry until cells reach the proper size ^45^. The particular control of quiescence and cell size in yeast makes it difficult to establish a parallelism with the function of MASTL in mammals. However, our data suggest that this kinase has evolved to participate in nutrient sensing by regulating the Ensa-PP2A/B55 pathway in the negative feedback loop that controls AKT-mTOR activity (Fig. S4d). Moreover, our data also suggest that mTORC1 can directly phosphorylate MASTL to regulate its activity in response to nutrients, another parallelism with the function of the yeast MASTL ortologue. MASTL appears to limit AKT activation in response to glucose and insulin but not in response to other nutrients, such as aminoacids, which are also sensed by mTORC1 pathway. Unfortunately, how glucose controls activation of the AKT-mTOR pathway is not completely understood ^46, 47^, and the observed selectivity of MASTL activity in glucose signaling deserves further studies. In *S. cerevisiae*, RIM15 activity and cellular localization is regulated by PKA and mTORC1 ^48-50^ and whether PKA control MASTL activity in mammals has not been explored yet.

In mammals, the insulin-mediated activation of AKT is critical for proper glucose disposal and metabolic adaptations. One of the key roles of AKT is the regulation of glucose uptake into insulin regulated tissues (reviewed in ^27, 51^. This is partially mediated by the translocation of GLUT4 from vesicular intracellular compartments to the plasma membrane. By phosphorylating and inhibiting glycogen synthase kinase (GSK3), AKT promotes glucose storage in the form of glycogen. In agreement with these data, increased AKT activity as a consequence of MASTL deletion leads to enhanced GLUT4 translocation and glucose uptake in cells, resulting in enhanced glycolysis (Fig 3). In vivo, deletion of Mastl leads to increase AKT phosphorylation and glycogen accumulation in liver, and enhanced GLUT4 translocation and glucose uptake in Mastl-deficient muscle cells (Fig. 4).

Insulin resistance is a pathological state of obesity and type 2 diabetes that occurs, at least in part, through chronic activation of mTORC1/S6K1 by nutrients and constitutive feedback activity that blocks AKT activation ^52^. Supporting the role of the mTORC1/S6K1 feedback loop in the pathogenesis of type 2 diabetes, S6K1-deficient mice are protected from age- and obesity-induced insulin resistance in the presence of reduced IRS1 phosphorylation ^53^. In addition, GRB10-deficient mice also exhibited enhanced glucose tolerance and insulin sensitivity ^54^, whereas IRS1-deficient mice developed insulin resistance ^55^. On the other hand, enhancing AKT signaling in skeletal muscle can improve systemic metabolism and overcome diet-induced obesity ^56^. Ablation of Mastl is also able to improve glucose tolerance in a model of high fat diet (HFD)-induced obesity and results in lower blood glucose levels in aged animals. The effect of Mastl ablation is likely mediated by increased PI3K/AKT signaling as it can be prevented upon treatment with a PI3K inhibitor. Of note, *Ensa* knockout mice have improved glucose tolerance and are more insulin sensitive ^57^. In the same direction, haploinsuficiency of the B55α subunit of PP2A causes insulin resistance likely due to the defective insulin-induced AKT stimulation in insulin-responsive tissues ^58^. These observations are in line with a model in which MASTL phosphorylates ENSA to inhibit PP2A/B55 and AKT signaling (Fig. S4d). These data suggest the intriguing possibility that MASTL inhibition could be considered as a new potential target of intervention for metabolic diseases, such as obesity-induced diabetes. Since MASTL does not participate in the central AKT-mTOR axis, its inhibition could be used as a more specific strategy to downregulate the negative feedback loop without the need of inhibiting the central mTOR-S6K kinases.

## Methods

### Cell lines

All human cancer cell lines were grown in DMEM supplemented with 10% FBS and gentamycin at 37°C with 5% CO2. MDA-MB231, BT549, and MCF7 breast cancer cell lines were kindly provided by M.A. Quintela (CNIO, Spain). HepG2 and C2C12 cells were kindly provided by A. Efeyan (CNIO, Spain) and P. Muñoz-Canoves (CNIC, Spain), respectively. Human embryonic kidney (HEK) 293-T cell line was obtained from American Type Culture Collection (ATCC). The MDA-MB231 cell line has been DNA fingerprinted using the GenePrint 10 System (Promega). All cell lines were routinely tested for mycoplasma.

### Genetically-modified mouse models

*Mastl* conditional knockout mice obtained were generated by crossing the *Mastl*(lox) allele with a mouse strain carrying a knock in of the CRE-ERT2 recombinase inducible by tamoxifen (TAM) in the locus of the ubiquitous RNA polymerase promoter (RNAP2_CreERT2)^21^. Males (8-12 week old) were fed for 9 weeks with 60% high fat diet (HFD) (60% fat, 20% protein, 20% carbohydrate; Research Diets; D12492). Mice were treated with TAM (Sigma; T5648) by intraperitoneal injection at a dose of 200 mg/kg twice one week before the assay.

All animals were maintained in a C57BL6/J-129S mixed background. Mice were housed in the pathogen-free animal facility of the CNIO (Madrid) and maintained under a standard 12-h light-dark cycle, at 23°C with free access to food and water. All animal work and procedures were approved by the ISCIII committee for animal care and research, and were performed in accordance with the CNIO Animal Care program following international recommendations.

### Plasmids, transfections and lentiviral infections

Silencing of *MASTL* and *TSC2* was performed using pLKO.1 lentiviral plasmids encoding specific siRNA (ON-TARGET SMARTpool, Dharmacon) or shRNA sequences (Sigma), as previously described ^59^. A shRNA scramble (SHC002, Sigma) was used as a control. *TSC2* was knocked out using the CRISPR-Cas9 system. The sgRNA against human *TSC2* gene ^60^ was subcloned into the lentiCRISPR v2 lentiviral vector (Addgene #52961). For lentiviral transduction, pLKO plasmids were co-transfected into HEK293-T cells along with packaging plasmids pMDLg/pRRE (Addgene #12251), pRSV-Rev (Addgene #12253) and pMD2.g (Addgene #12259) expression plasmids using Lipofectamine 2000 (Invitrogen) following manufacturer’s instructions. Virus-containing supernatants were collected 24 and 48 hours after transfection and target cells were infected in the presence of 4μg/mL polybrene. Cells were analyzed 64 h later, and knockdown efficiencies were evaluated by immunoblotting using specific antibodies. For conditionally knockout *MASTL* gene in the MDA-MB231 cell line, we used a doxycycline-inducible CRISPR-Cas9 system ^59^. *isgMASTL* clones constitutively express the sgRNA targeting the exon 1 of *MASTL*, and Cas9 expression is induced by doxycycline (Dox) at a concentration of 2 μg/mL. Experiments were performed 64-72 h after Dox addition. To knock down *PPP2R2A* and *PPP2R2D* transcripts, specific siRNAs (SI02225825 for *PPP2R2A* and SI02759148 for *PPP2R2D*)) were purchased from Qiagen. ON-TARGET plus SMARTpool siRNAS against human *ENSA* (L-011852-00) and *ARPP19* (L-015338-00) were purchased from Dharmacon. A siRNA against luciferase was used as a control (Dharmacon). Transfections were performed using Hiperfect (Qiagen), according to manufacturer’s instructions.

For transient overexpression, the full-length cDNA of mouse Mastl was amplified by PCR using mouse cDNA as a template and subcloned into the pDEST 3.1-GFP vector (Gateway system, Invitrogen). Kinase-dead mutation (G43S) and phosphorylation mutants (S861A, S864A) were generated by site-directed mutagenesis. For rescue assays, full-length human cDNAs for ENSA, and ARPP19 were amplified by PCR using cDNA clones (MGC:839; IMAGE:2820528 for ENSA and MGC:5468; IMAGE:3451558 for ARPP19) and subcloned as N-terminal Flag- and HA-tag fusions, respectively, into the lentiviral vector pLVX.puro (Clontech). Phosphomimetic mutations (ENSA-S67D, ARPP19-S62D) were generated by site-directed mutagenesis. Lentivirus production was performed as indicated above, and host cells were infected with viral supernatants and selected with puromycin (1 μg/ml).

### Nutrient deprivation assays

For nutrient-deprivation experiments, cells were seeded into 6 well plates at a density of 300,000 cells per well 24 h before starvation. Actively proliferating cells were rinsed twice with Hank’s balanced salt solution (HBSS) and incubated in DMEM without glucose (Gibco) or DMEM/Ham’s F-12 without amino acids (US Biological) supplemented with 10% dialyzed FBS (dFBS, Gibco) for 1 h, unless otherwise indicated. Stimulation with 25 mM glucose (Sigma) was performed for 10, 15 or 60 min, as indicated. For serum withdrawal, cells were rinsed twice with HBSS and incubated in medium containing 0.1% FBS for 6 h and, when specified, stimulated with 100 nM insulin (Sigma). For oxygen starvation, cells were treated with the hypoxia-mimetic agent 150 μM CoCl_2_ (Sigma) for 22 h in complete media. Incubation with inhibitors was performed at the time of starvation or 10-20 min before re-stimulation. For mTOR inhibition cells were treated with rapamycin (Santa Cruz Biotechnologies) or torin1 (Selleckem) at 100 and 250 nM, respectively. For PP2A inhibition okadaic acid (Calbiochem) and fostriecin (Santa Cruz Biotechnologies) were used at 50 nM and 5 μM, respectively.

### Immunodetection

For immunoblot analysis, cells were extracted on ice in cold lysis buffer (50 mM Tris-HCl pH 7.5, 1 mM EDTA, 1 mM EGTA, 0.1% Triton X-100, 0.1% 2-mercaptoethanol) in the presence of protease inhibitors (Roche) and phosphatase inhibitors (1 mM sodium orthovanadate, 50 mM sodium fluoride, 20 mM β-glycerophosphate). Mouse tissues were extracted in RIPA buffer (50 mM Tris-HCl pH 7.5, 150 mM NaCl, 1% NP-40, 0.5% Sodium Deoxycholate, 0.1% SDS) with protease and phosphatase inhibitors using a Precellys tissue homogenizer. Lysates were cleared were cleared by centrifugation at 15,000 g for 10 min at 4°C, and protein was quantified using the Bradford method. Protein extracts were mixed with sample buffer (350 mM Tris-HCl pH 6.8, 30% glycerol, 10% SDS, 0.6 M DTT, 0.1% bromophenol blue), boiled for 5 min and separated on TGX Criterion Bis-Tris acrylamide gels (BioRad). Proteins were then transferred to a nitrocellulose membrane (BioRad), blocked in PBS 0.1% Tween-20 buffer containing 3% non-fat dried milk, and probed overnight with primary antibodies (1:1000 dilution) at 4 °C and for 1 h at room temperature with peroxidase-conjugated secondary antibodies. Blots were developed using enhanced chemiluminiscence reagent (Western Lightning Plus-ECL; Perkin Elmer), exposed to an autoradiograph film and developed using standard methods. Films were scanned and quantifications were performed using ImageJ.

Antibodies for western blot (WB) were obtained from the following sources: anti phospho-T308 AKT (9275), phospho-S473 AKT (9271), phospho-T1462 TSC2 (3611), phospho-S939 TSC2 (3615), phospho-T389 S6K1 (9234), phospho-S235/236 S6 (2211), phospho-S612 IRS1 (3203), phospho-S307 IRS1 (2381), phospho-S476 GRB10 (11817), phospho-T202/Y204 ERK (9101), phospho-S79 ACC (11818), phospho-S792 Raptor (2083), phospho-T246 PRAS40 (2640), phospho-S9 GSK-3β (9336), phospho-S67/S62 Ensa/Arpp19 (5240), total AKT (9272), ERK (9102) TSC2 (4308), S6K1 (9202), S6 (2217), IRS1 (3407), GRB10 (3702), ACC (3676), Raptor (2280), PTEN (9188) were from Cell Signaling Technology. Anti-human and mouse MASTL (MABT372), and phospho-S10 H3 (06-570) were from Millipore. Anti-mouse Mastl (AP14289c) was from Abgent and anti-human MASTL was generated at the CNIO (available from Abcam, ab166647). Anti-pan-B55 (sc-81606) was from Santa Cruz. Anti phospho-T194 MASTL was a gift from H. Hochegger (Univ. of Sussex). Anti ENSA (180513), and HA (1424) were from Abcam. Anti β-Actin (5441), FLAG (F7425) and Vinculin (V9131) were from Sigma. Anti GFP was from Roche (11814460001). Mouse monoclonal anti-GAPDH was generated at the Monoclonal Antibodies Unit at the CNIO.

For immunofluorescence, MDA-MB231 cells cultured on glass coverslips were fixed in 4% buffered formaldehyde (FA) for 10 min at RT. For GLUT4 immunodetection, FA was first quenched with 100 mM glycine for 10 min followed by permeabilization using 0.2 % Triton X-100 for 10 min, and blocking with 10% goat serum plus 1% BSA for 1 h at RT. Incubation with a specific antibody against GLUT4 (ab654, 1:200) was performed for 1 h at 37°C in combination with 1 μM HCS cell mask (Invitrogen; H32720) that stains the entire cell, and 10 ng/ml of 4’,6-Diamidine-2’-phenylindole dihydrochloride (DAPI) to visualize the nuclei.

For histological examination, samples were fixed in a solution of 4% of paraformaldehyde, embedded in paraffin and cut in 2.5-μm sections. Then, sections were deparaffinised in xylene and rehydrated through a series of graded alcohols, stained with hematoxylin and eosin (H&E) and Periodic-Schiff staining (PAS staining), following standard protocols. Additional immunohistochemical examination was performed using a specific antibody against Ki67 (Master Diagnostica). For immunofluorescence, rehydrated paraffin sections were boiled for 5 min in 10 mM citrate buffer at pH 6.0 for antigen retrieval and then they were incubated 1 h at room temperature with normal goat serum blocking solution (2% goat serum, 1% BSA, 0.1% cold fish skin gelatine, 0.1% Triton X-100, 0.05% Tween-20 and 0.05% sodium azide in PBS at pH 7.2). Sections were incubated overnight at 4 °C with the indicated primary antibodies: Glut4 (ab654, Abcam, 1:100), Phospho-IRS1 (Ser616) (44-550G, Thermofisher, 1:100); Vinculin (V9131, Sigma, 1:200). Slides were then washed three times for 3 min in 0.05% of Tween-20 in PBS and incubated with Alexa-conjugated secondary antibodies (Molecular Probes) for 1 h at room temperature in 3% BSA in PBS. After washing three times for 3 min in PBS, cell nuclei were counterstained with 100 ng/ml of DAPI for 10 min at room temperature (Roche Applied Science). Slides were mounted on coverslips using Fluoromount mounting medium (DAKO). Images were captured using a laser scanning confocal microscope TCS-SP5 (AOBS) Leica. Image analysis was performed using ImageJ software.

### In vitro kinase assays

Cells transiently expressing GFP-mMastl were lysed in ELB buffer (50 mM Hepes pH 7.4, 150 mM NaCl, 5 mM EDTA, 1% NP-40) in the presence of protease and phosphatase inhibitors. Cell lysates were cleared by centrifugation at 15,000 g for 10 min at 4°C. Mastl immunocomplexes were purified with an anti-GFP antibody (Roche), crosslinked to Dynabeads Protein A (Invitrogen). Immunoprecipitates were washed two times in ELB buffer, twice in high salt (500 mM NaCl) ELB buffer, followed by two additional washes in kinase buffer (20 mM Hepes pH 7.5, 10 mM MgCl_2_). Equal amounts of beads were re-suspended in kinase buffer and mixed with 500 ng of recombinant ARPP19 purified from HEK293T cells (kindly provided by A. Castro, Montpellier, France) in a final volume of 20 μL, in the presence of 50 μM ATP and 2 μCi ^32^P-labeled ATP, and incubated for 20 min at 30°C. Reactions were stopped by the addition of sample buffer and boiling for 5 min. Samples were analyzed by SDS-PAGE in a 10% acrylamide-gel, transferred to a nitrocellulose membrane, and developed by exposure to a radiosensitive screen. Images and quantifications were performed using ImageJ. Total MASTL in the reactions was quantified by immunoblot against MASTL on the transferred membrane.

For mTORC1 and p70S6K1 kinase assays, active recombinant proteins were purchased from Sigma (SRP0364, P0066, respectively). Recombinant purified human MASTL protein was used as a substrate and was from Abnova (H00084930). Reaction was performed in a final volume of 30 μL consisting of kinase buffer (supplemented with 50 mM KCl for the mTOR assay), 200 ng of the upstream kinase and 500 ng of MASTL, in the presence of 100 μM ATP and 2 μCi ^32^P-labeled ATP, and incubated for 30 min at 30°C. Reactions were stopped by the addition of sample buffer and boiling for 5 min, and were analyzed by SDS-PAGE in an 8% acrylamide-gel. Proteins in the gel were stained using Imperial Stain (Invitrogen) following the manufacturer’s instructions. The gel was vacuum-dried and exposed to a radiosensitive screen.

### Mastl phosphorylation site mapping

mTORC1 and S6K1 in vitro kinase assays were performed as detailed above but in the absence of radiolabeled ATP. Samples in 5 M urea were doubly digested using the FASP procedure with some modifications. The resulting peptides were analyzed by LC-MS/MS using an Impact mass spectrometer (Bruker Daltonics). Raw files were searched against a Uniprot *Homo sapiens* database (20,187 sequences) using Andromeda as the search engine through the MaxQuant software. Peptide identifications were filtered by Percolator at 1% FDR using the target-decoy strategy. Label-free quantification was performed and extracted ion chromatograms for MASTL phosphopeptides were manually validated.

To map phosphoresidues on Mastl, endogenous mouse Mastl was immunoprecipitated from mouse embryonic fibroblasts (MEFs) using a polyclonal antibody against Mastl (Abgent; AP14289c) crosslinked to Dynabeads Protein A. Cells were extracted in RIPA buffer as indicated above and samples were subjected to mass spectrometry analysis. Protein eluates were doubly digested using the FASP procedure with some modifications. The resulting peptides were analyzed by LC-MS/MS using a LTQ Orbitrap Velos mass spectrometer (Thermo Scientific). In addition, samples were subjected to TiO2 enrichment. Raw files were searched against a UniprotKB/TrEMBL *Mus musculus* database (UniProtKB/Swiss-Prot/TrEMBL 43,539 sequences) using Sequest-HT as the search engine through the Proteome Discoverer 1.4 (Thermo Scientific) software. Peptide identifications were filtered by Percolator at 1% FDR using the target-decoy strategy. Label-free quantification was performed with MaxQuant and extracted ion chromatograms for MASTL phosphopeptides were manually validated in Xcalibur 2.2 (Thermo Scientific).

### Metabolite analysis by NMR

Concentrations of cell media metabolites were determined by NMR spectroscopy as described in ^61^. Briefly, HepG2 cells were transduced with scramble or shMASTL lentiviruses. 48h post-infection cells were plated in DMEM containing 10 mM U-^13^C-glucose and 10% FBS, and samples were taken at variable times after incubation in fresh medium. Glucose and lactate concentrations were measured from the sum of all resolved signals in the first increment of a 2D ^1^H-^1^H NOESY experiment, using the H4 signal of glucose and the methyl signal of lactate. Concentrations were normalized to the number of cells in each condition.

### Glucose uptake assays

Cells were serum-deprived for 16h, washed twice in KRH buffer (50 mM Hepes pH 7.4, 137 mM NaCl, 4.7 mM KCl, 1.85 mM CaCl_2_, 1.3 mM MgSO_4_ and 0.1% (w/v) BSA), and incubated for 5 min at 37°C in KRH buffer with 6.5 mM 2-deoxy-D-[1-^3^H]-Glucose (0.5 μCi) and 7 mM D-[^14^C]-mannitol (0.3 μCi) per well. Then, cells were washed four times with ice-cold KRH buffer and lysed with 0.05M NaOH for 2h at 37°C. Cell lysates were mixed with 3 ml of liquid scintillation cocktail and radioactivity was quantified using a Perkin Elmer *Wallac 1414* Liquid Scintillation counter. From the ^14^C specific radioactivity the extracellular water volume present in the culture was calculated. This enables the extracellular ^3^H volume to be determined and subtracted from the total ^3^H radioactivity to calculate the intracellular amount of 2-deoxy-D-[1-^3^H]-Glucose transported.

Ex vivo muscle [^3^H]-2-deoxy-D-glucose uptake was assessed using methods previously described, with minor modifications^62, 63^. Briefly, Soleus and Extensor Digitorum Longus (EDL) muscles from both legs were dissected out from overnight fasted and anesthetized mice with ketamine (100 mg/kg) and xylacine (10 mg/kg). Mice were euthanized by cervical dislocation after muscles had been removed and muscles were rapidly incubated for 60 min at 37 °C in pre-gassed (95% O_2_ and 5% CO_2_) Krebs-Ringer bicarbonate (KRB) solution containing the following (in mmol/L): 117 NaCl, 4.7 KCl, 2.5 CaCl_2_·2H_2_O, 1.2 KH_2_PO_4_, 1.2 MgSO_4_·7H_2_O, and 24.6 NaHCO_3_ and supplemented with 2 mmol/L pyruvate and 0.1% of BSA (RIA Grade). For insulin stimulation, muscles were incubated in the presence or absence of insulin (100 nM; Sigma-) for 20 min at 37 °C. Contralateral non-stimulated muscles were used as controls. Glucose transport was assessed during 10 min at 37 °C using 1 mM 2-Deoxyglucose (2-DG) containing 1.5 μCi/ml 2-deoxy-D-[^1, 2-3^H]-glucose (Perkin Elmer, Waltham, MA) and 7 mM mannitol containing 0.45 μCi/ml D-[^14^C]-mannitol (Perkin Elmer, Waltham, MA) in KRB with or without 100 nM insulin. The gas phase was maintained at 95% O_2_ and 5% CO_2_. After radioactive incubations, muscles were dried by putting them briefly on Whatmann paper, flash-frozen in liquid nitrogen, weighted, and solubilized for 30 min at 65 °C in 300 μl 1 M NaOH. Non-soluble particulates were precipitated by centrifugation at 10,000 xg for 1 min and aliquots were removed for quantification of [^3^H] and [^14^C] labels in a liquid scintillator counter. Presence of [^14^C]-mannitol was used to correct for the amount of extracellular [^3^H]-2-DG that was non-specifically uptaken by the tissue but not actively transported into muscle cells. Radioactive counts were normalized to muscle weight.

### Metabolic assays in mouse models

For glucose tolerance test assay (GTT) mice were fasted for 16 h and then administered an intraperitoneal glucose injection (2 g/kg body weight). Glycaemia was measured by tail vein blood sampling using a glucometer and glucose strips (Bio Medical) in fasting and 15, 30, 45, 60, 90 and 120 min after glucose injection. The PI3K inhibitor CNIO ETP-44692 ^28^ was administered to the 16 h fasted mice by oral gavage in a unique dose of 20 mg/kg 1 h before starting the GTT. ETP-44692 was dissolved in 90% PEG-300 and 10% N-methyl-2-pyrrolidone (NMP). Insulin tolerance test (ITT) was performed in mice fasted for 6 h and then intraperitoneally injected with rapid acting insulin (0.75 U/kg body weight) (Actrapid, Novo Nordisk) following by tail vein blood glucose sampling to measure glycaemia at 0, 15, 30, 45 and 60 min after insulin injection.

### Statistics

Statistical analysis was carried out using Prism 6 (GraphPad). Statistical tests were performed using two-sided, unpaired Student’s *t*-test, paired Student’s *t-*test, or 1-or 2-way ANOVA (Bonferroni’s multiple test) or Chi-square test according to specifications in the figures. Data with p>0.05 were considered not statistically significant (ns); *, p<0.05; **, p<0.01; ***, p<0.001.

## Acknowledgements

We are fully indebted to B. Manning (Harvard TH Chan School of Public Health, Boston) and A. Efeyan (CNIO) for suggestions and helpful discussions. We thank H. Hochegger (University of Sussex) for antibodies, A. Castro (University of Montepellier) for recombinant proteins, A.Efeyan (CNIO), M.A. Quintela (CNIO) and P.Muñoz-Canoves (CNIC and Pompeu Fabra University) for cell lines, and Dario Hermida (CNIO) for help with 3D structural modeling. B.S.C. was supported by Foundation la Caixa, and A.E.B. was funded by Comunidad de Madrid. M.A.F. was supported by a young investigator grant from the Spanish Ministry of Economy and Competitiveness (MINECO; SAF2014-60442-JIN; co-financed by FEDER funds). The Cell Division and Cancer lab of the CNIO is supported by grants from the MICIU (RTI2018-095582-B-I00), Red de Excelencia iDIFFER (RED2018-102723-T), Comunidad de Madrid (B2017/BMD-3884), and Worldwide Cancer Research (WCR-20-0155).

## Author Contribution

B.S.-C., B.H., A.E.B., B.S., D.M.-A., D.H. and J.G-M. carried out the experiments and analyzed the data. C.S. and R.C.-O. performed and analyzed the NMR studies, and P.X. and J.M. carried out the mass spectrometry studies. M.A.-F. and M.M. designed the project, analyzed the data and wrote the manuscript.

