## Supplementary Figures for "A cell cycle kinase-phosphatase module restrains PI3K-Akt activity in an mTORC1-dependent manner"

By Belén Sanz-Castillo et al.

### Supplementary Figures

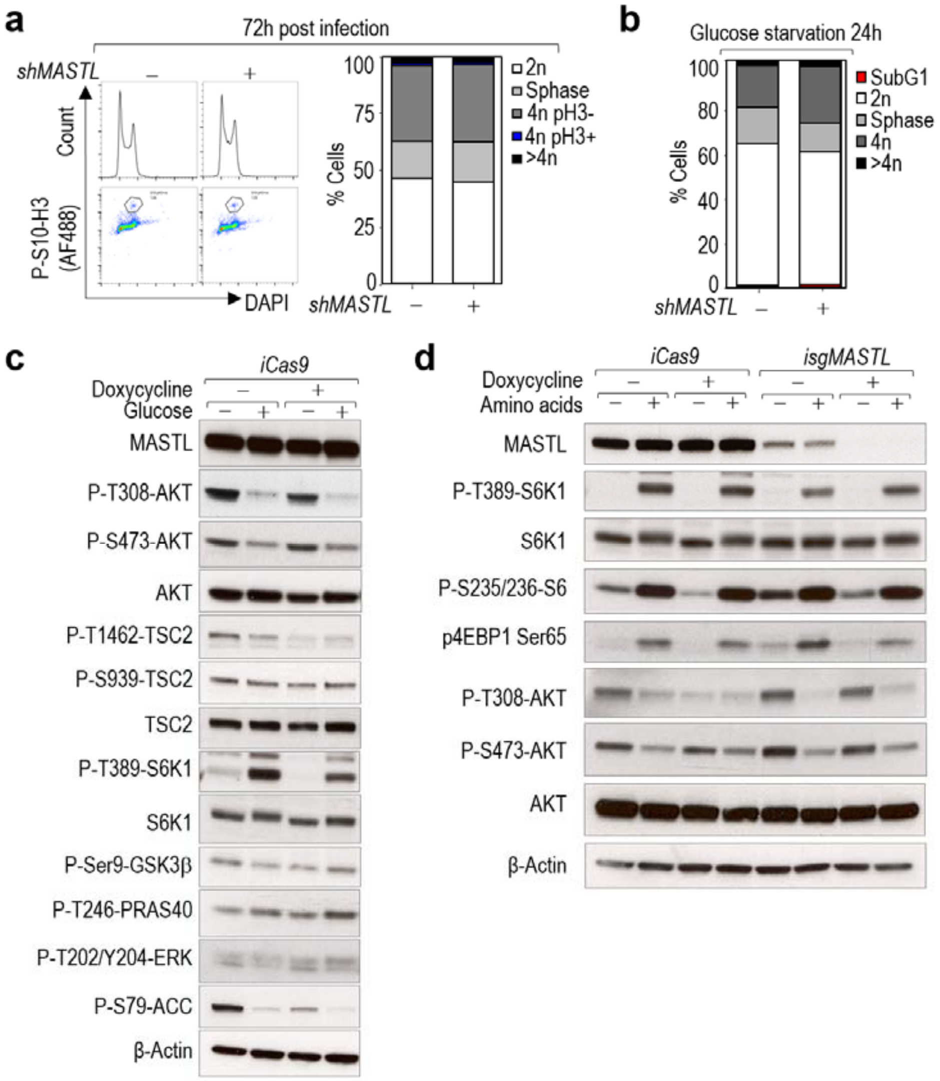


Supplementary Figure 1 | MASTL modulates AKT activity in response to glucose independently of its mitotic function (Related to Fig. 1). a) Quantification of the different phases of the cell cycle in control and *MASTL* null cells. MDA-MB-231 cells were infected with shRNAs specific for *MASTL* (+) or Scramble (–) as a control, and cell cycle was analyzed 72 h later. Mitotic cells were scored using immunodetection of P-S10-H3 (pH3) and total DNA was stained with DAPI followed by flow cytometry analysis. Left pictures show representative cell cycle profiles and P-S10-H3 staining. b) Quantification of the percentage of cells in the different phases of the cell cycle before or 24 h after glucose starvation. Cell cycle analysis was performed by flow cytometry. DNA content was determined by propidium iodide staining. c) Control of the Dox and Cas9 expression in the MDA-MB-231 *isgMASTL* system using a control clone that expresses Cas9 alone (iCas9). Same protocol as in Fig. 1e was followed. d) CRISPR/Cas9-mediated ablation of *MASTL* using the CRISPR/Cas9 inducible system (*isgMASTL* and *iCas9* as a control). 72 h after Dox, cells were maintained in complete medium (+) or starved of aminoacids for 1 h. Whole-cell lysates were blotted with the indicated antibodies.

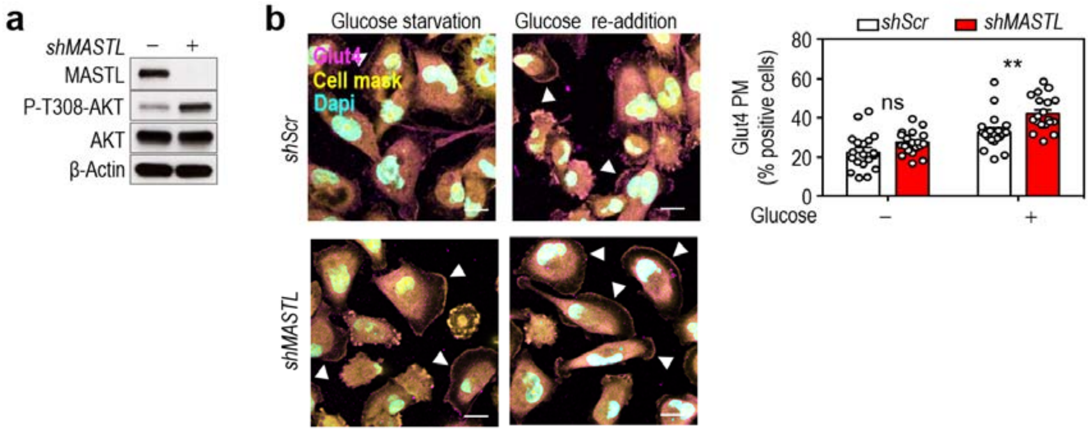


Supplementary Figure 2 | Effect of MASTL depletion on cellular metabolism (Related to Fig. 3). a) Immunoblot confirming *MASTL* depletion and activation of the AKT pathway in *MASTL*-null cells in the conditions of the assay shown in Fig. 3a and Supplementary Fig. S2b). MASTL was knocked down using shRNAs for *MASTL* (+) and Scramble (–) sequences as a control. b) Immunofluorescence for GLUT4 in MDA-MB-231 cells in conditions of glucose starvation and re-addition. Representative micrographs of GLUT4 (magenta) are shown. Nuclei were stained with DAPI (cyan) and cell mask is shown in yellow. Arrowheads indicate positive cells for GLUT4 translocation to the plasma membrane (PM; right panel) in 18 fields/condition. Plots represent quantification of the percentage of cells positive for GLUT4 translocation to the PM (Bottom panel). At least 800 cells were counted per condition. Error bars indicate s.d. Ns, not significant; **p<0.005; ***p<0.001 (2-way ANOVA).

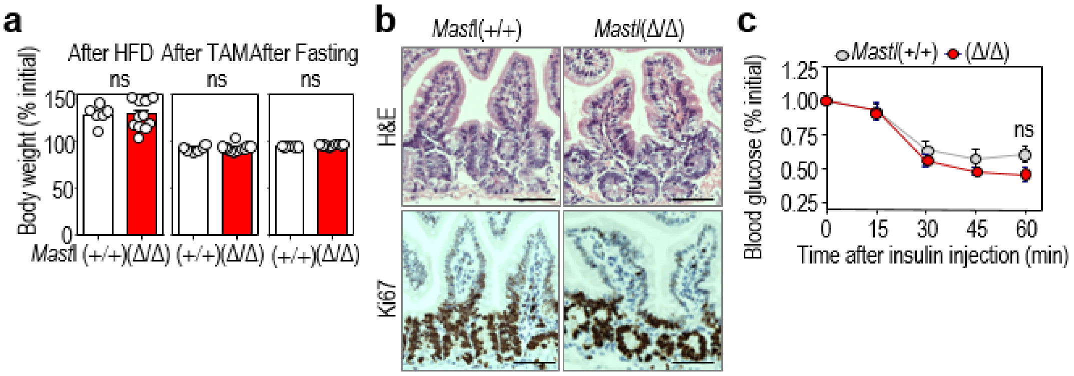


Supplementary Figure 3 | Effect of MASTL ablation on glucose metabolism in vivo (Related to Fig. 4). a) Plots representing the relative body weight of the mice included in the GTT assay from Fig. 4. Body weight gain after 9 weeks of HFD (Left); Body weight loss 1 week after tamoxifen injection (Middle); Body weight loss after 16 h fasting (Right). n=6 *Mastl*(+/+) and n=11 *Mastl*(Δ/Δ). Error bars indicate s.e.m. (two-sided student's *t*-test). b) Representative images of the intestine from *Mastl*(+/+) and *Mastl*(Δ/Δ) mouse showing normal architecture of the epithelia by hematoxylin and eosin (H&E) staining (upper panel) and similar levels of proliferation (Ki67 staining, lower panels). c) Insulin tolerance test (ITT) in *Mastl*(+/+) (n=6) and *Mastl*(Δ/Δ) (n=11) mice. Data are mean ± s.e.m. ns, not significant; 2-way ANOVA.

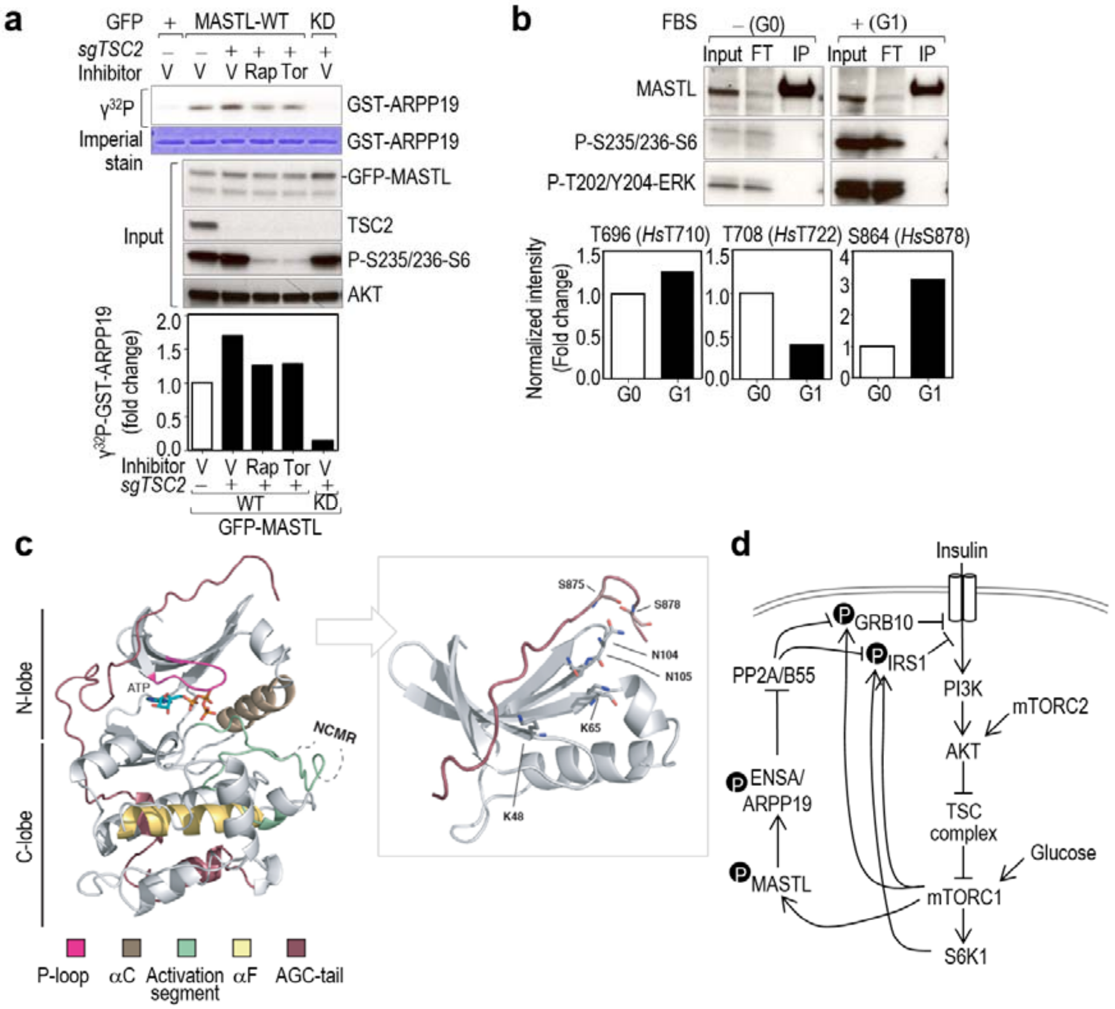


Supplementary Figure 4 | Regulation of MASTL activity by mTORC1 (Related to Fig. 7). a) *In vitro* kinase assay using GFP-hMASTL immunocomplexes from asynchronous TSC2 knock out cells (*sgTSC2*) or control cells treated with the indicated inhibitors for 1 h. Immunoprecipitation of GFP alone was used as a control. GST-ARPP19 was used as a substrate, and autophosphorylation of MASTL was used as a readout of MASTL activity. Upper panel shows the radiograph of ARPP19 phosphorylation in the different conditions accompanied by the immunoblots of the whole cell extracts. Quantification of the relative kinase activity of MASTL is shown in the lower graph, in which MASTL activity in control cells was set as 1. b) Mouse embryonic fibroblasts (MEFs) were synchronized in G0 by serum starvation and then released for 6h in the presence of serum (G1). Endogenous Mastl was immunoprecipitated with a specific antibody against Mastl (Abgent) and subjected to mass-spectrometry to identify phosphorylated residues in Mastl in the indicated conditions. Upper panel shows the immunoblot of the immunoprecipitates and inputs probed with the indicated antibodies. Lower graphs show the relative quantification of the indicated phosphopeptides in both conditions. c) A homology structural model showing the position of S875 and S878 in the regulatory C-tail domain of human MASTL. Ribbon diagram of the MASTL kinase domain (without the residues 181-727 coding for the NCMR; left panel) in complex with ATP. The protein fold displays the standard bilobal structure (N-lobe and C-lobe) with its respective AGC-tail (shown in brown). Please see the associated key for common structural kinase domain components. Molecular details for the phospho-binding pocket of MASTL (right panel). Using the homology-modeled AGC C-tail of MASTL, it is possible to determine that amino acids Lys48, Lys65, Asn104 and Asn105 (stick representation) are positioned such that they could interact with the phosphorylated tail/liker reside (pSer875) and the mTOR-phosphorylation site (pSer878). Note, in both panels, the MASTL protein has been homology-modeled using MODELLER based on the structures of MASTL (PDB 5LOH2; https://www.rcsb.org/) for the kinase domain and of PKA and RSK2 (PDBs 1ATP and 4EL9, respectively) for the AGC C-tail. **d)** A model for the role of the MASTL-ENSA/ARPP19-PP2A/B55 pathway in the negative feedback loop that controls AKT activity in response to mTORC1-S6K1 signaling. For simplicity, only the phospho-residues in the negative feedback loop are shown.
